# Determining controllability of sepsis using genetic algorithms on a proxy agent-based model of systemic inflammation

**DOI:** 10.1101/153080

**Authors:** Chase Cockrell, Gary An

## Abstract

Sepsis, a manifestation of the body’s inflammatory response to injury and infection, has a mortality rate of between 28%-50% and affects approximately 1 million patients annually in the United States. Currently, there are no therapies targeting the cellular/molecular processes driving sepsis that have demonstrated the ability to control this disease process in the clinical setting. We propose that this is in great part due to the considerable heterogeneity of the clinical trajectories that constitute clinical “sepsis,” and that determining how this system can be controlled back into a state of health requires the application of concepts drawn from the field of dynamical systems. In this work, we consider the human immune system to be a random dynamical system, and investigate its potential controllability using an agent-based model of the innate immune response (the Innate Immune Response ABM or IIRABM) as a surrogate, proxy system. Simulation experiments with the IIRABM provide an explanation as to why single/limited cytokine perturbations at a single, or small number of, time points is unlikely to significantly improve the mortality rate of sepsis. We then use genetic algorithms (GA) to explore and characterize multi-targeted control strategies for the random dynamical immune system that guide it from a persistent, non-recovering inflammatory state (functionally equivalent to the clinical states of systemic inflammatory response syndrome (SIRS) or sepsis) to a state of health. We train the GA on a single parameter set with multiple stochastic replicates, and show that while the calculated results show good generalizability, more advanced strategies are needed to achieve the goal of adaptive personalized medicine. This work evaluating the extent of interventions needed to control a simplified surrogate model of sepsis provides insight into the scope of the clinical challenge, and can serve as a guide on the path towards true “precision control” of sepsis.

**Author summary:** Sepsis, characterized by the body’s inflammatory response to injury and infection, has a mortality rate of between 28%-50% and affects approximately 1 million patients annually in the United States. Currently, there are no therapies targeting the cellular/molecular processes driving sepsis that have demonstrated the ability to control this disease process. In this work, we utilize a computational model of the human immune response to infectious injury to offer an explanation as to why previously attempted treatment strategies are inadequate and why the current approach to drug/therapy-development is inadequate. We then use evolutionary computation algorithms to explore drug-intervention space using this same computational model. This allows us to characterize the scale and scope of interventions needed to successfully control sepsis, as well as the types of data needed to derive these interventions. We demonstrate that multi-point and time-dependent varying controls are necessary and able to control the cytokine network dynamics of the immune system.

## Introduction

Approximately 1 million people will be diagnosed with sepsis, a condition with a mortality rate ranging from 28%-50%, each year [1]. Attempts to discover biologically-targeted therapies for sepsis have thus far been focused on manipulating a single mediator/cytokine, generally administered with either a single dose or over a very short course (<72 hours) [2,3]. Unfortunately, all these attempts, have been unsuccessful [2,3], likely due to both the nonlinear nature of the human inflammatory signaling network and the paucity of clinical time-course data to place network relationships in context. In fact, we would propose that the universal failure to find effective cellular/molecular control strategies effective at the clinical level raises the question as to whether the system can be effectively controlled at all. The rationale for the current investigation is an attempt to address this fundamental question: can the trajectory of clinical sepsis be controlled, and if so, what is the scale and scope of the therapeutic interventions required to do so?

It is well known in biology that the systemic response to identical perturbations in genetically identical individuals (i.e., mice) is governed according to some probability distribution. In a chaotic system, this small stochastic variability in response can ultimately lead to a radically different final state [4]. It logically follows then that a single time point/single cytokine intervention is unlikely to be successful on a broad range of patients with a broad range of conditions that have led to the state of sepsis. The challenge, however, is that the range of possible interventions, which is a function of the number of potential molecular targets, the extent to which they are modified, the time at which such modification can occur and the combinations thereof, is staggering, and cannot be tractably investigated given the logistical and practical limitations of both experimental and clinical research. We propose to address this challenge by the use of evolutionary computing (in the form of genetic algorithms) applied to a sufficiently complex, albeit abstracted, proxy computational model of sepsis.

We have previously proposed that dynamic computational modeling, and specifically agent based modeling, can be used to represent mechanistic biological knowledge in a way that reproduces the non-linear dynamics of the real world system [5,6]. ABM’s have been used to study and model a wide variety of biological systems [7], from general purpose anatomic/cell-for-cell representations of organ systems capable of reproducing multiple independent phenomena [6] to platforms for drug development [8], and are frequently used to model non-linear dynamical systems such as the human immune system [9,10]. Specifically, was have previously developed an agent-based model (ABM) of systemic inflammation, the Innate Immune Response agent-based model (IIRABM). We propose to use the existing IIRABM as a surrogate proxy system [11] for the investigation of potential control strategies for sepsis. We note that while the model does not contain a comprehensive list of all signaling mediators present in the human body, all relevant cellular behaviors are represented. The named cytokines in this model are those that are most often associated with the available behavior rules in the current literature [12]. The IIRABM is a two-dimensional abstract representation of the human endothelial-blood interface. This abstraction is designed to model the endothelial-blood interface for a traumatic (in the medical sense) injury, and does so by representing this interface as the unwrapped internal vascular surface of an azimuthally averaged 2D projection of the terminus for a branch of the arterial vascular network. This abstraction serves two primary purposes: to allow circumferential access to the traumatic injury by the innate immune system, and to incorporate multiple levels if interaction between leukocytes and tissue. The IIRABM utilizes this abstraction to simulate the human inflammatory signaling network response to injury; the model has been calibrated such that it reproduces the general clinical trajectories of sepsis (see supplemental material for details). The IIRABM operates by simulating multiple cell types and their interactions, including endothelial cells, macrophages, neutrophils, TH0, TH1, and TH2 cells as well as their associated precursor cells. The simulated system dies when total damage (defined as aggregate endothelial cell damage) exceeds 80%; this threshold represents the ability of current medical technologies to keep patients alive (i.e., through organ support machines) in conditions that previously would have been lethal. The IIRABM is initiated using 5 external variables – initial injury size, microbial invasiveness, microbial toxigenesis, environmental toxicity, and host resilience. In previous work [13], we have performed a parameter sweep over these variables to determine the plausible boundaries for conditions that could be considered clinically relevant. These are parameter sets that can lead to multiple outcomes – complete healing, death by infection, or death from immune dysregulation/sepsis.

Additionally, the IIRABM has been used to perform in silico clinical trials of mediator-directed therapies via the inhibition or augmentation of single and specific cytokine synthesis pathways [12]. Those studies accurately reproduced unsuccessful clinical trials, as well as the non-efficacy of hypothetical interventions; however to date no effective putative interventions have been discovered.

The human innate immune response, and specifically in terms of sepsis, can be characterized through measurement of various biomarkers, including the pro-inflammatory and anti-inflammatory cytokines included in the IIRABM [14]. This implies that the system can be characterized by its state at some specific time, and therefore we apply that same perspective to our investigations with the IIRABM. At each time step, the IIRABM measures the total amount of cytokine present for all mediators in the model across the entire simulation. The ordered set of these cytokine measurement creates a high-dimensional trajectory through cytokine space that lasts throughout the duration of the simulation (until the in silico patient heals completely or dies). Prior analysis of these trajectories has shown that the aggregate output of the IIRABM behaves as a Random Dynamical System (RDS) with chaotic features [13] (in the sense that future simulation state can be sensitive to initial conditions). A Random Dynamical System [15] can be described by the triplet (*S*, Γ, *Q*) where S is the state space, Γ is a family of maps which maps *S* back onto itself (often referred to as the “equations of motion”), and *Q* is some probability distribution on Γ. Simply put, an RDS is a system in which the equations of motion (in this case, the equations which give the aggregate cytokine value for the system at a specific instance in time) contain elements of randomness. A detailed discussion of this, and more formal definition, can be found in [16].

System state in the IIRABM can be defined by a vector of cytokines, 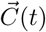, in which each element 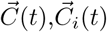, is the total amount of an individual cytokine present in the simulation at that instance in time. The cytokines which comprise this vector are: Platelet-activating factor (PAF), Tumor necrosis factor alpha (TNF*α*), Soluble tumor necrosis factor receptors (sTNFr), Interleukin-1 (IL1), soluble interleukin-1 receptors (sIL1r), Interleukin-1 receptor antagonist (IL1ra), Interferon-gamma IFN*γ*, Interleukin-4 (IL4), Interleukin-8 (IL8), Interleukin-10 (IL10), Interleukin-12 (IL12), and Granulocyte colony-stimulating factor (GCSF). At each time step, 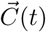 is given by:

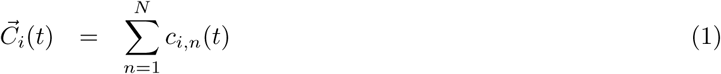

where *c_i,n_*(*t*) is the individual cytokine concentration seen by the endothelial cell at a specific grid point and *N* is the total number of endothelial cells which make up the endothelial surface. The random element comes from the calculation:
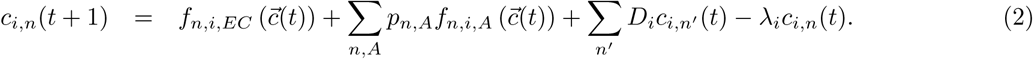

This equation contains four primary terms: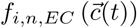 is the amount of cytokine *i* produced by endothelial cell *n* in response to current cytokine concentrations; 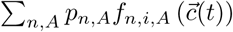 is the sum of the responses of all other cell types, indexed with *A*, at the location of endothelial cell *n*, where *p_n,A_* is the population of a specific cell type at that location; 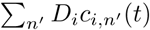 represents the amount of cytokine *i* which diffuses into location *n* (according to the associated diffusion constant, *D_i_*) from the neighboring cell locations, denoted by *n’;* and *λ_i_c_i_*_,*n*′_(*t*) is the amount of cytokine at cell location *n* that degrades at each time step according to degradation constant *λ_i_.* The randomness in this model comes from the term *p_n,A_.* This population is stochastically random in two ways: 1) in the absence of cytokine markers, inflammatory cell movement follows a random walk, and 2) cellular differentiation of inflammatory cells proceeds according to a probability distribution parameterized by current cytokine concentrations. We note that this explicit randomness does not comprise the entire uncertainty present in the model. The aggregation of individual cytokine concentrations into a single measure conceals any spatial information present in the simulation. Due to this, future behaviors can appear to be random when, in reality, an epistemic uncertainty (as opposed to intrinsic or “real” stochasticity randomness) prevents an accurate prognosis. A high-level overview of this system and visual depiction of the functional rules governing cytokine production can be seen in Fig. 1.

**Fig 1.**
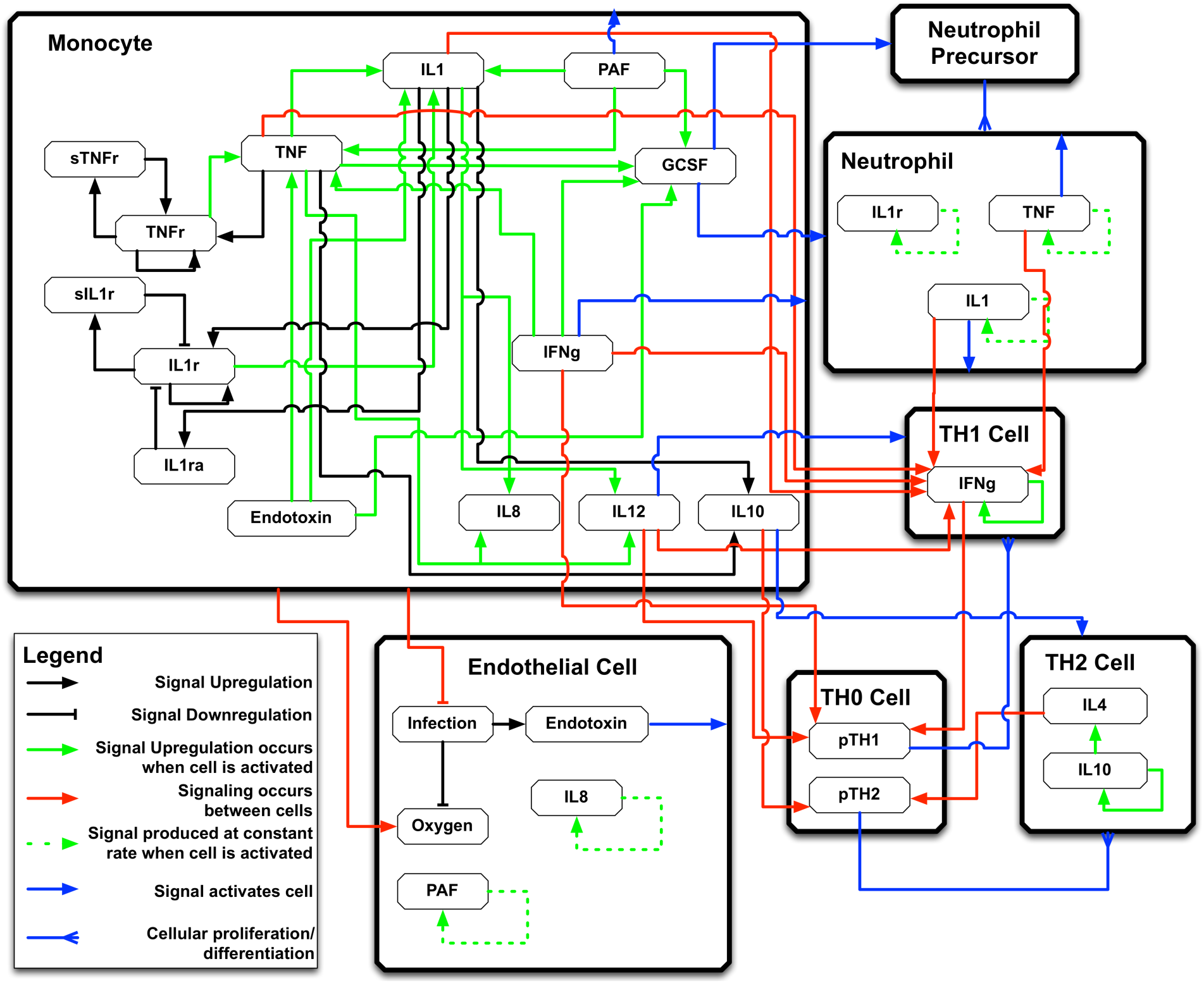
A high-level overview of the signaling rules incorporated into the IIRABM. Rules represented include cytokine up-regulation/down-regulation, cell activation, and cellular differentiation.

Discovery of an effective or optimal intervention can then be viewed as a nonlinear optimization of a control problem [17–19]. Genetic Algorithms (GA) [20] and Evolutionary Computing have been used to optimize a wide variety [21] of nonlinear systems. Medical applications of GA include vaccine dosing strategies and protein binding site prediction [22,23]. Given a sufficiently validated model, and interpretation of the development of an intervention/control strategy to be a nonlinear optimization problem, GA’s can also be utilized to develop complex treatment and control strategies.

## Results

The parameter set (invasiveness=2, toxigenesis=5, host resilience=0.1, environmental toxicity=2) upon which the GA was trained generated a set of simulations that led to a final outcome of death by sepsis with a probability of 82%, therefore representing a challenging case to digitally “cure.” Table 1 displays results from a variety of interventions that minimized their associated fitness function functions.

**Table 1.** Mortality rates (MR) for varying numbers and lengths of independent interventions.

| Number of Interventions | Length (hrs) | General MR | Patient Specific MR |
| --- | --- | --- | --- |
| 1 | 3 | 80 | 65 |
| 1 | 6 | 80 | 64 |
| 1 | 12 | 95 | 87 |
| 2 | 3 | 77 | 67 |
| 2 | 6 | 73 | 66 |
| 2 | 12 | 75 | 71 |
| 3 | 6 | 56 | 41 |
| 3 | 12 | 44 | 30 |
| 6 | 12 | 25 | 12 |
| 7 | 12 | 30 | 12 |
| 8 | 12 | 16 | 12 |

The first column displays the number of sequential interventions (spaced 12 hours apart); the second column displays the length of the intervention in simulation time steps; the third column displays the general probability of death for this specific condition (parameter set) after intervention; the fourth column displays the probability of death for the specific in silico patient upon which the GA was trained. The general probability of death was calculated by starting the simulation with 100 distinct RNG seeds, applying the selected intervention, and running the simulation to completion. The probability of death for the specific patient was calculated by starting the simulation with a specific RNG seed, reseeding the RNG 100 times at the start of the intervention sequence, applying the intervention, and running the simulation to completion (either complete healing or death). We have previously characterized the stochastic properties of the trajectory space of the IIRABM in [13], and note the existence 3 distinct regions of state space: 1) the region of space under the influence of the Life Attractor – trajectories in this region will always lead to a state of complete healing; 2) the region of space under the influence of the Death attractor – trajectories in this region will always lead to complete system failure and death; and 3) the “stochastic zone” – trajectories in this region are influenced more strongly by random effects (stochastic randomness and epistemic uncertainty) than by the effects of either of the aforementioned attractors.

At the time point when the intervention is started, the system is still in the “stochastic zone,” and thus a given intervention will not have a guaranteed effect; rather, it changes the probabilistic future outcome distribution to favor a state corresponding to greater health. Thus, a subtle effect sustained over time causes a significant improvement in mortality rates. Throughout the course of this work, we initiate our interventions 12 hours post-simulated injury. The GA has found an effective intervention which begins at this point, however, if stochasticity leads an in silico patient to a sufficiently different location in cytokine state-space, the stochastic failure of the first stage of the intervention will lead to further failure at subsequent stages. The nature of these dynamics explains the phenomenon of “non-responders” to the putative interventions as defined at various time points during an individual trajectory; ultimately “non-responders” represent trajectories that are either not driven out of the stochastic zone by our proscribed duration of therapy or those whose stochastic response lead them to a region of parameter space in which the intervention no longer alters the future outcome probability distribution to favor a state of increased health.

The best solution (that which minimized the probability of death for both the specific patient case and the general case) was found by using 8 sequential interventions. For the specific patient upon which the GA was trained, the probability of death was reduced from 68% to 12% through application in the intervention shown in Fig 2; for the general case, the probability of death with this intervention was reduced from 82% to 16%. This intervention is represented as a three-dimensional bar graph in Fig 2. The height of the bars along the z-axis represents the log2 of the intervention multiplier; the x-axis enumerates the interventions; the y-axis shows which cytokine has its protein synthesis augmented or inhibited according to its associated bar.

**Fig 2.**
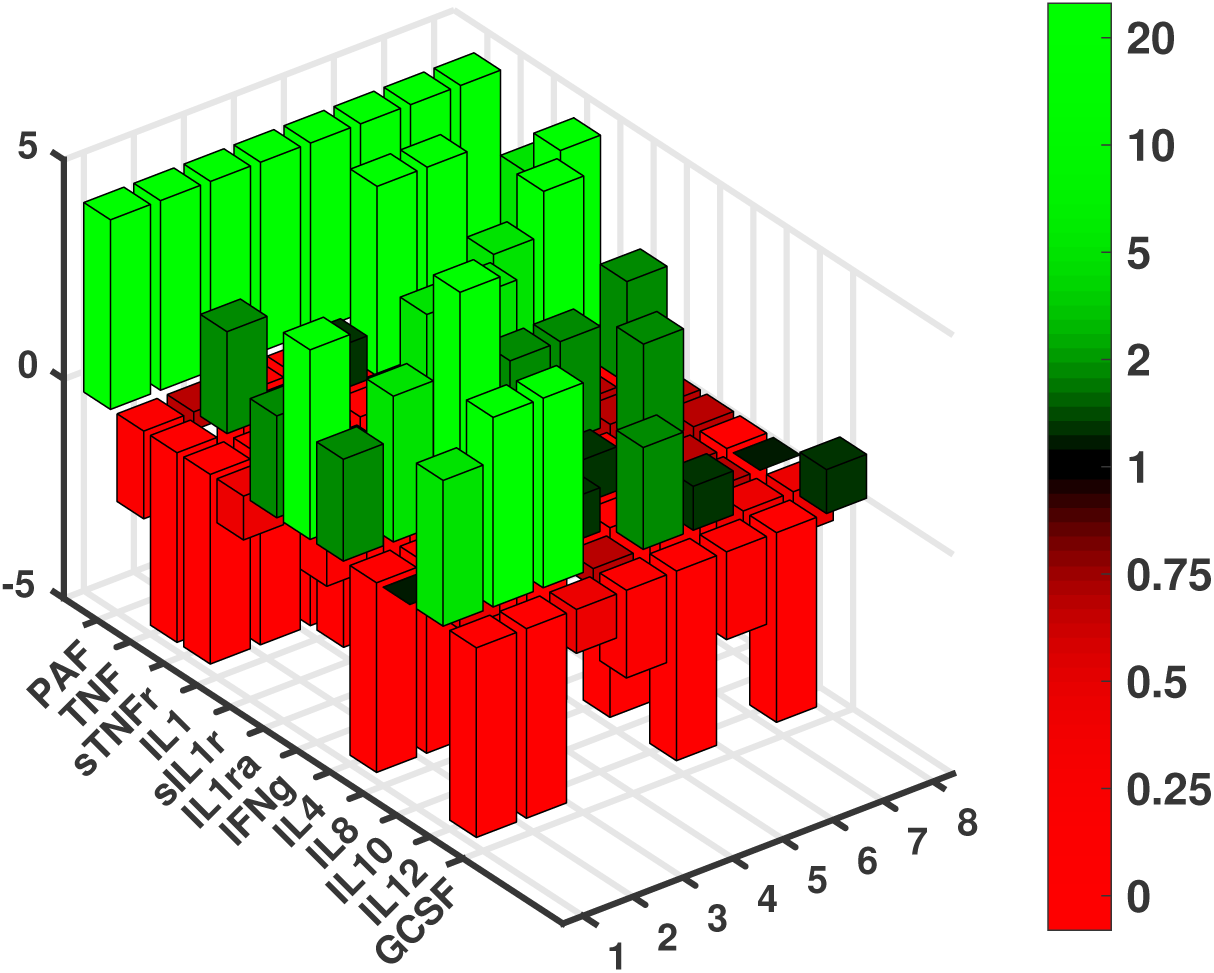
A 3d bar graph representing the series of 8 sequential interventions to which the genetic algorithm (GA) converged. The sequential order is displayed on the x-axis; the cytokines operated on are displayed on the y-axis; the base-2 logarithm of the augmentation or inhibition strength is displayed on the z-axis.

In order to explore the generalizability of this intervention, we tested it against all parameter sets that generated a mortality rate between 1 and 99% - this is the population of parameter sets that have been defined as “clinically relevant” [13]. The results of this test are shown in Fig. 3. In Panel A, we show the untreated mortality rate distribution for all parameter sets that generate at least two outcomes (life and death). This distribution is skewed towards the edge cases as the number of combinations that generates multiple outcomes is finite, while those that generate a single outcome are unbound; i.e., if a host resilience parameter value of 5 generates healing 100% of the time, all parameter values greater than 5 will lead to healing 100% of the time [13]. Panel 3B shows the overall shift in the mortality rate distribution for this population of parameter sets. Fig. 3, Panels C-F show the shifts in mortality rate distribution for more narrow ranges of mortality. We note that Fig. 3, panels E and F appear to have a bimodal distribution (also hinted at in Panel D); this bimodal distribution occurs because as the severity of the simulated condition increases, the likelihood of a given stochastic replicate being a “non-responder” to a standardized treatment also increases.

**Fig 3.**
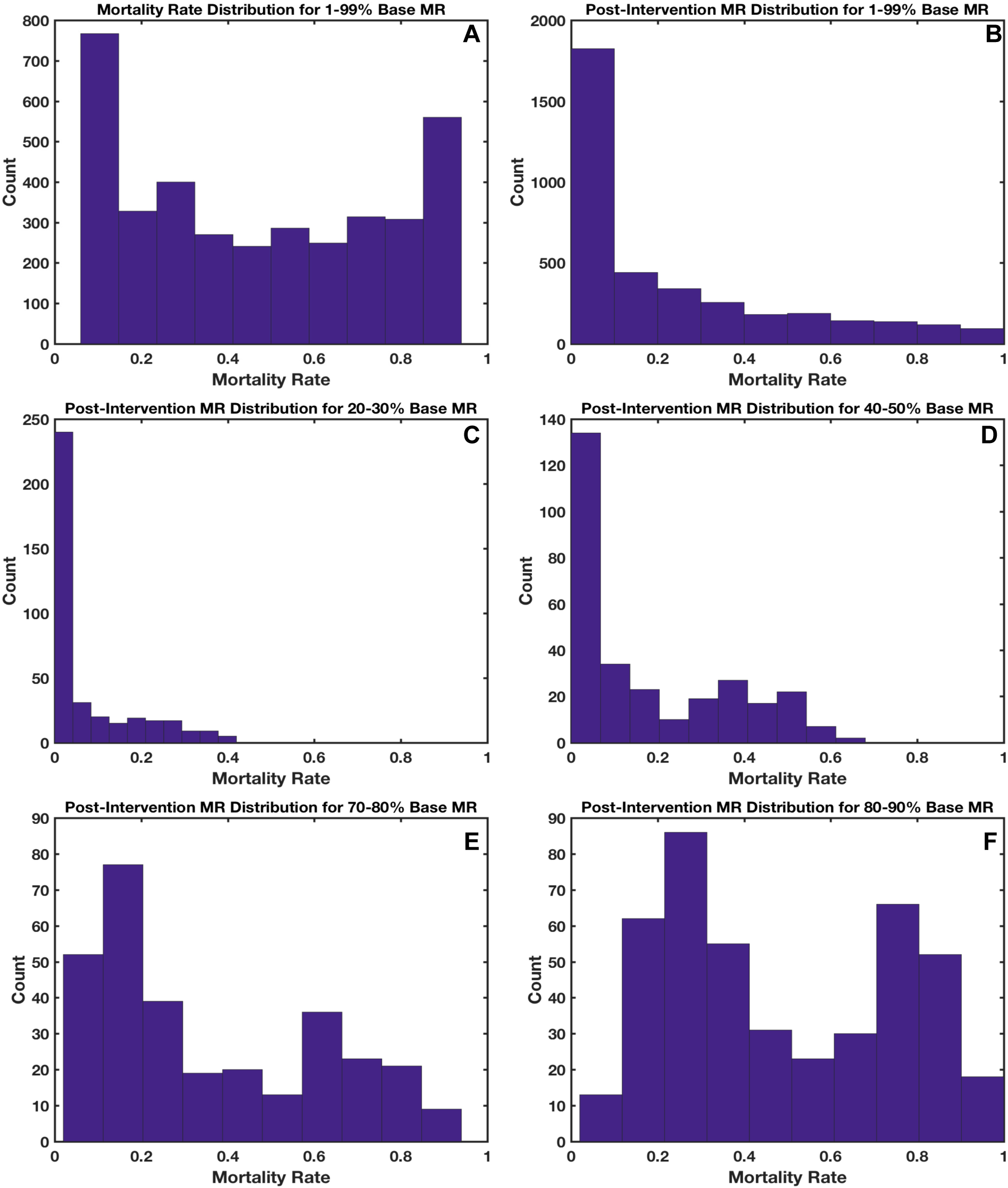
In panel A, the mortality rate distribution for the parameter sets that generate between 1 and 99% mortality rate (MR) is presented with the mortality rate on the x-axis and the number of parameter sets that generate that mortality rate (with 100 stochastic replicates) is on the y-axis. Panel B shows the MR distribution for the same set of simulations as in panel A, however they have been treated with the calculated intervention. Panel C shows the post intervention MR for those simulations in panel A which have a base MR between 20% and 30%; Panel D shows the post intervention MR for those simulations in panel A which have a base MR between 40% and 50%; Panel E shows the post intervention MR for those simulations in panel A which have a base MR between 70% and 80%; Panel F shows the post intervention MR for those simulations in panel A which have a base MR between 80% and 90%.

The phenomenon of “non-responders” provides an excellent argument for *adaptive personalized medicine,* that is, the need to adapt a therapeutic strategy based on an individual’s response to therapy (“in silico clinical trials of 1”), with the goal of returning a system to a state of health. To further explore this, we investigated the cases that were unable to be cured using the derived intervention. Fig. 4A shows the total oxygen deficit trajectories for the average of all the simulations that healed when using the optimal intervention and for a single instance of the simulation that does not heal; Fig 4B shows the neutrophil population and Fig 4C shows the total systemic GCSF. After three sequential interventions, it is apparent that this patient has a stronger response to GCSF stimulation, and thus a higher neutrophil population. As this difference continues to compound over time, interventions later in the sequence lose their efficacy (as they are optimized for a system in a different state). After the application of three sequential interventions, it is apparent that this particular *in silico* patient is not responding desirable to the therapy. We pause the simulation at this point and use the GA to find a sequence of 5 more interventions that could heal the system. The newly derived final 5 interventions are compared with the old intervention in Fig. 5. This shifted the probability of death for that patient (at the time in which the intervention is changed) from 75% to 8%. This success suggests that an algorithm that adapts interventions based on system response could be more successful than a GA.

**Fig 4.**
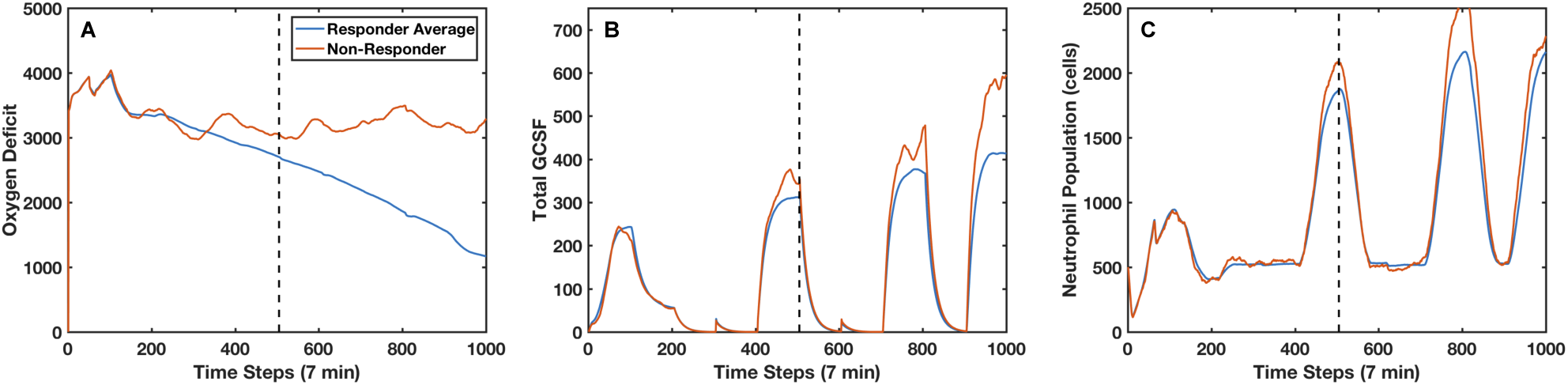
Panel A displays the oxygen deficit (an inverse measure of the system’s health) for an intervention non-responder (red) compared to the average oxygen deficit for intervention responders (blue) over time. Panel B displays the total GCSF for the non-responder and the responder average; panel C displays the total neutrophil population for an individual non-responder and the responder average. In this case, the non-responder does not end up healing due to a hyper-productive response to GCSF pathway stimulation, which leads to a surplus neutrophil population; this patient ultimately dies due to inflammation, which is exacerbated by the applied intervention.

**Fig 5.**
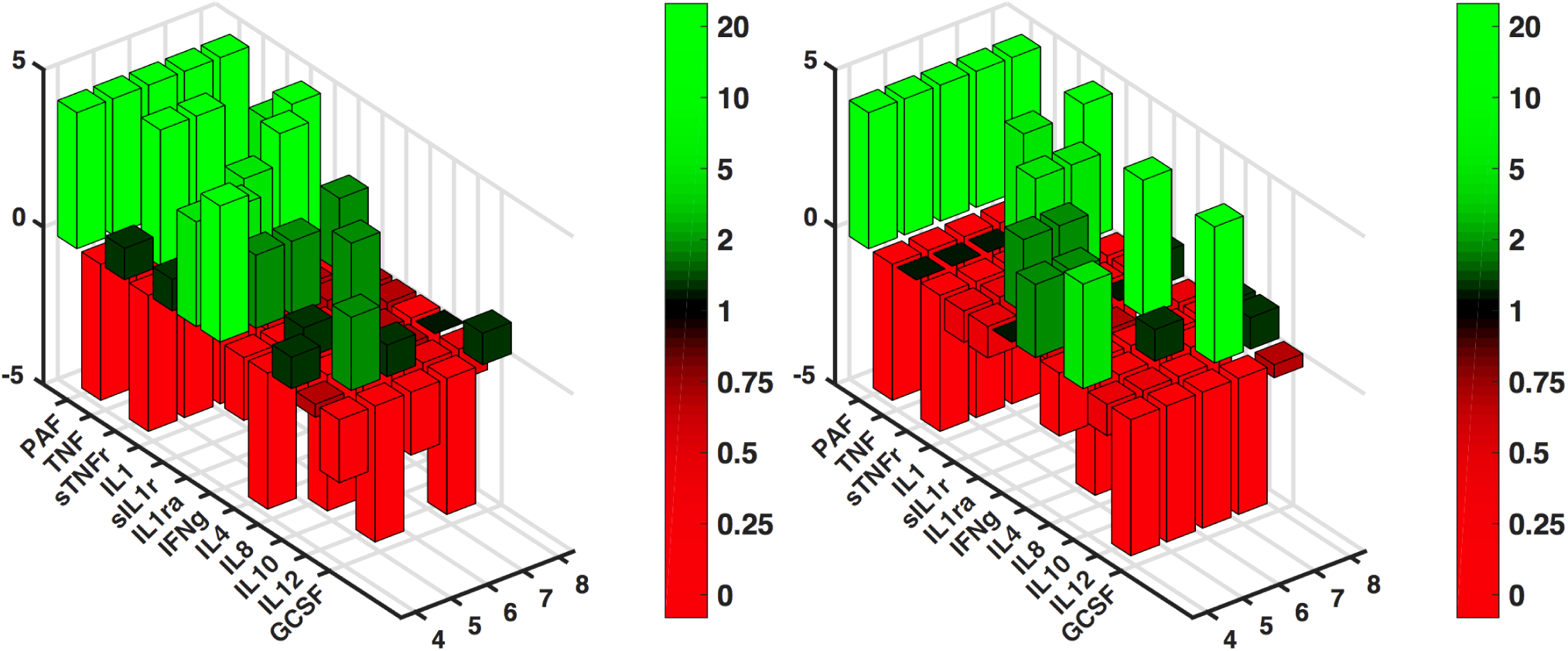
The bar graph on the left shows the final 5 interventions in a sequence of 8 which showed the greatest success in healing the most in silico patients. *Note that the first three intervention steps are identical to those shown in Fig. 2.* The bar graph on the right shows an alternate intervention sequence that was generated by training the genetic algorithm on a patient who was non-responsive to the original intervention. The in silico patients received identical interventions for the first three time points. At time point3, a significant deviation from expected behavior is noted in the non-responder. At this time point, the simulation is halted and used to populate the sample set for a new run of the genetic algorithm. When given the original intervention, this patient has a 75% chance of death at 60 hrs post-injury; the alternate intervention lowers this chance to 8%.

## Discussion

The human inflammatory signaling network as represented by the IIRABM is nonlinear, stochastic, and chaotic system that lacks a unique set of analytical equations that can adequately describe it. Designing an effective strategy to control such a system requires the exploration of an astronomically large search space – in the case of 8 sequential interventions with 9 defined augmentation/inhibition strengths, there are approximately 1091 combinations that can be applied to the system. Given the size of this space, we make no claim that we have found the globally optimal intervention for our model, given a fixed set of parameters; rather, we have shown that GA can be used to find a “good-enough” solution that shows success (though not perfection) at treating a range conditions leading to death by sepsis.

Longer duration therapies were more successful than shorter therapies due to the stochastic nature of the IIRABM and its response to perturbations (see supplemental material). All interventions are performed on in silico patients whose system state is either in the stochastic zone or under the influence of the death attractor (in which case their trajectories would have to pass through the stochastic zone on the way to health). When the system state is in the stochastic zone, the effects of randomness can be stronger than the effects of either of the attractors which influence the system; subsequent states will fall within some probability distribution (which is discoverable via pausing the simulation and reseeding the random number generator several times) based on the current state. In that sense, we could also consider the aggregate cytokine and health output to be a Markov Random Field [24]. Just as the system evolves in a stochastic nature when left unattended, its responses to perturbations will also be stochastic, however the possible future distribution of states is now based on the components of the perturbation as well as the current system state. Successful interventions will then shift the future state probability distribution towards health; more successful interventions will generate a larger move towards health more often, though there can still exist a non-zero probability that the system state evolves negatively rather than beneficially.

While GA is quite successful at healing at IIRABM under a wide range of conditions, it has a few drawbacks which preclude it from being the ideal solution: 1) more extreme conditions (either very large injuries or extremely virulent bacteria) require either a finer degree of control than is computationally tractable using GA, as each sequential intervention multiplies the size of the search space by a factor of 912 (approximately 5 billion), or more aggressive interventions; intervention multipliers were limited to a small set of values we considered clinically tractable – removing this constraint would lead to an unconstrained search, increasing computational cost and potentially generating implausible interventions; 2) adjusting the temporal density of interventions requires a new run of the GA, which can be computationally expensive; 3) the GA does not have the ability to react to non-responders and adjust the intervention accordingly – rather, it finds the single sequence of interventions which (locally) maximizes the survival probability for a given patient population. We should note that we do not claim to find the absolute optimum sequence of interventions for a given parameter set due to the lack of general formal analytical convergence proofs for genetic algorithms [25,26] and the fact that it is computationally intractable to explore the entire intervention space, especially for multiple sequential interventions. Additionally, many interventions can have opposing effects with the possibility of cancelling each other out (i.e., GCSF augmentation leading to an increasing neutrophil population).

Future work will incorporate alternative machine learning algorithms, including deep reinforcement learning [27] and Long Short Term Memory neural networks [28] for time-series prediction of aggregate cytokine values. Both of these techniques would base putative interventions on the sequence of events that lead up to the intervention. In this sense, the learning algorithm would adapt the putative intervention to an individual system state rather than attempting to develop a single broad policy that would cover a certain class of injury.

These results strengthen the case for adaptive personalized medicine, a therapeutic strategy which adapts and evolves in real time based on patient response. Rather than searching an astronomically large space for a utilitarian intervention, personalized medicine techniques would respond to cytokine dynamics with an individualized intervention for each patient at varying temporal scales. These things are theoretically possible using GA, but at the present time, the computational expense limits the practicality of using this technique to personalize treatments.

## Materials and methods

The current investigation involves providing a proof-of-concept example of identifying whether effective controls can be found for the IIRABM, and, by extension, for the treatment of sepsis. This initial proof-of-concept constrains the problem by focusing a particular parameter configuration of the IIRABM with a mortality rate of 68%. We first employed a genetic algorithm (GA) to search for possible therapeutic interventions, and then examined the generalizability of the optimal solutions across a wider range of stochastic replicates and additional parameter sets with similar overall baseline mortality rates. The IIRABM was implemented in C++ and simulations were performed on the Edison Cray XC30 Supercomputer at the National Energy Research Scientific Computing Center and on the Beagle Cray XE6 Supercomputer at the University of Chicago.

We chose to train the GA on a single parameter set for 2 primary reasons: 1) A large number of parameter sets can generate a realistic sepsis condition. Parameter sets that are relatively similar tend to have similar mortality rates and similar simulated length-of-stays in the ICU. In this work, we explored the generalizability of intervention solutions derived using the GA to help assess the future potential of utilizing GA as an in silico drug-development technique. 2) The computational cost of running a genetic algorithm is substantial. A single instance of this model simulating 90 days in the ICU will take approximately 4 minutes (depending on the speed of the processor running the simulation). Each iteration of the GA gathers data from up to 2000 independent simulation runs (where each run repeats the simulation 10 times using 10 stochastic replicates), and the GA can run for 1000 or more generations before convergence when considering multi-stage interventions; each GA experiment can utilize up to 1,000,000 cpu-hrs (equivalent to $100,000) [29].

We have selected a parameter set (invasiveness=2, toxigenesis=5, host resilience=0.1, environmental toxicity=2) which leads to simulated death approximately 80% of the time in the general case. This is illustrated in Figure 6A; here we show total health trajectories for 100 stochastic replicates of the above parameter set. The systemic oxygen deficit (a proxy for total system damage) is plotted against the time for which the simulation ran. Trajectories shown in red are simulations that die and trajectories shown in blue are simulations that heal completely. From this set of outcome trajectories, we have chosen a specific trajectory/in silico patient with which to train the GA. Given this specific patient’s location in state space 24 hours post injury, we estimate that they have a 68% chance of death. Note that this mortality rate is calculated in a different manner than the rate referenced above. In the general case, this parameter set generates a mortality rate of 80%, meaning that when the simulation’s random number generator (RNG) is given 100 unique starting seeds, 80 of those simulations will die (a population level outcome distribution). For the individual patient mortality rate, the simulation is paused at the time step before the intervention would be applied (in this case, 24 hrs post injury); the simulation’s RNG is then reseeded a number of times to discover the mortality rate for a specific in silico patient at a specific moment in time. Thus, the general case mortality rate represents the mortality rate for a specific parameter set, while the individual patient mortality rate represents the mortality rate for a specific parameter set at a specific instance in time. As discussed above, an individual patient does not have a fixed fate at any given point in time (as long as their trajectory remains in the stochastic zone) and their probabilistic future outcome distribution is time dependent and will evolve with the system. This is shown in Figure 6B; this image displays 100 random number generator (RNG) re-seedings, reseeded 24 hours post injury, of the specific trajectory we have chosen. The utilization of stochastic replicates on a single patient allows the GA to sample the full range of responses possible for a given intervention.

**Fig 6.**
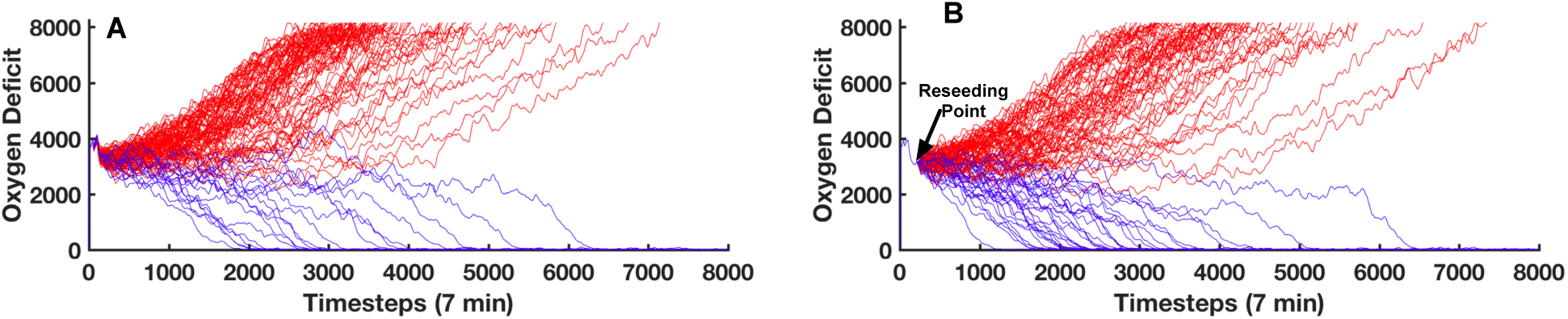
Panel A displays 100 stochastic trajectories generated by the IIRABM for a specific parameter set (Invasiveness=2; Host Resilience=0.1; Toxigenesis=5; Environmental Toxicity=2) with an injury with a radius of 33 cells. The total systemic oxygen deficit, an inverse measure of the in silico patient’s health (y-axis) is presented as a function of time (x-axis). 82% of the simulated patients (red) end in death, while 18% heal completely (blue). The trajectories shown in panel B use the same parameter set as in panel A, however the run is started with a specific random number generator seed; the random number generator is re-seeded at 1 day post injury. At 1-day post injury, this specific patient has a 68% chance of death.

We have selected a set of cytokines and associated targets (PAF, TNF*α*, sTNFr, IL1, sIL1r, IL1ra, IFN*γ*, IL4, IL8, IL10, IL12, and GCSF), which are the principal drivers of the inflammatory/immune dynamics expressed by the model. In order to search for an optimal intervention strategy, we allow production of each of these targets to be augmented or inhibited alone or as a group. For the purposes of this study, we consider an intervention to be a set of signaling augmentations/inhibitions. An individual’s chromosome is then a 1x12n vector, where n is the number of independent sequential interventions, containing this information. The augmentation and inhibition values are selected from the set: 0.05, 0.25, 0.5, 0.75, 1, 2, 5, 10, 20; inhibitory values are multiplicative (i.e., *S_i,new_* = *S_i_*_,*old*_ *\* I_i_*) where *S_i,new_* represents the modified signal value, *S_i,old_* represents the signal value being modified, and *I_i_* represents the vector element of the intervention vector which applies to the signal.

Augmentations are additive rather than multiplicative (i.e., *S_i,new_* = *S_i,old_* + (*I_i_*_−1_). This allows for a sustained application of the intervention; were the augmentations to be multiplicative, then the simulation would quickly produce results outside the realm of plausibility due to an exponential explosion in concentrations of cytokines whose values have been multiplied. To illustrate this, consider *S*_0_ = 10; we wish to intervene on this cytokine’s production for 100 time steps by augmenting production by a value of 2 (20%) – if we use the additive method of augmentation, on the 100th time step, 200 units of cytokine have been added. Alternatively, using a multiplicative method, a 20% increase would be a multiplication by a factor of 1.2 – after 100 time steps, this cytokine has increased by a factor of 1.2100, or approximately 80 million.

We define the fitness function as 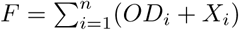, where *OD* represents the Oxygen Deficit, and is an inverse measure of the patient’s total health at the time when the fitness is evaluated and *X* is a measure of the total infectious load in the system at the time when the fitness is evaluated, and n is the number of stochastic replicates used. An optimal solution will minimize this fitness function (and thus maximize the patient’s total health). We train the GA on 10 stochastic replicates of an individual trajectory. These replicates are generated by re-seeding the RNG at the time point when the intervention begins. This allows the GA to learn from a possible range of responses and individual can have to a given intervention. The fitness is evaluated 12 hours after the application of the final intervention (which has a maximum duration of 12 hours).

After the fitness is evaluated, we use the tournament selection method [30,31] with a tournament size of 2 to generate the breeding population. When breeding, each pair of progenitors produces two progeny using a uniform crossover operator [32]. We continue this process until the algorithm has converged to a small set of possible solutions, and select the solution that leads to the minimal fitness value as the intervention to be tested. We evaluate the intervention according to three criteria: 1) outcome distribution in specific patient upon which the GA was trained; 2) outcome distribution in a population of patients exposed to the same injury and microbial infection; and 3) outcome distribution in a population of patients exposed to a range of injuries and microbial infections.

Source code for the IIRABM with GA capability is available at https://bitbucket.org/cockrell/iirabm public. Certain GA experiments used the EMEWS [33] framework with DEAP [34] to automate the GA process.

## Acknowledgments

This research used high performance computing resources of the National Energy Research Scientific Computing Center, a DOE Office of Science User Facility supported by the Office of Science of the U.S. Department of Energy under Contract No. DE-AC02-05CH11231. Additionally, this research was supported in part by NIH through resources provided by the University of Chicago Computation Institute (Beagle2) and the Biological Sciences Division of the University of Chicago and Argonne National Laboratory, under grant 1S10OD018495-01. This work was also supported by funds from Lawrence Livermore National Laboratory under Award #B616283.

